# Broad anti-coronaviral activity of FDA approved drugs against SARS-CoV-2 *in vitro* and SARS-CoV *in vivo*

**DOI:** 10.1101/2020.03.25.008482

**Authors:** Stuart Weston, Christopher M. Coleman, Rob Haupt, James Logue, Krystal Matthews, Matthew B. Frieman

## Abstract

SARS-CoV-2 emerged in China at the end of 2019 and has rapidly become a pandemic with roughly 2.7 million recorded COVID-19 cases and greater than 189,000 recorded deaths by April 23rd, 2020 (www.WHO.org). There are no FDA approved antivirals or vaccines for any coronavirus, including SARS-CoV-2. Current treatments for COVID-19 are limited to supportive therapies and off-label use of FDA approved drugs. Rapid development and human testing of potential antivirals is greatly needed. A quick way to test compounds with potential antiviral activity is through drug repurposing. Numerous drugs are already approved for human use and subsequently there is a good understanding of their safety profiles and potential side effects, making them easier to fast-track to clinical studies in COVID-19 patients. Here, we present data on the antiviral activity of 20 FDA approved drugs against SARS-CoV-2 that also inhibit SARS-CoV and MERS-CoV. We found that 17 of these inhibit SARS-CoV-2 at a range of IC50 values at non-cytotoxic concentrations. We directly follow up with seven of these to demonstrate all are capable of inhibiting infectious SARS-CoV-2 production. Moreover, we have evaluated two of these, chloroquine and chlorpromazine, *in vivo* using a mouse-adapted SARS-CoV model and found both drugs protect mice from clinical disease.

## Introduction

At the end of December 2019, reports started to emerge from China of patients suffering from pneumonia of unknown etiology. By early January, a new coronavirus had been identified and determined as the cause (1). Since then, the virus originally known as novel coronavirus 2019 (nCoV-2019), now severe acute respiratory syndrome coronavirus 2 (SARS-CoV-2), has spread around the world. As of April 23^th^, 2020, there have been roughly 2.7 million confirmed cases of COVID-19 (the disease caused by SARS-CoV-2 infection) with over to 189,000 recorded deaths (www.WHO.org). Multiple countries have enacted social distancing and quarantine measures, attempting to reduce person-to-person transmission of the virus. Healthcare providers lack pharmaceutical countermeasures against SARS-CoV-2, beyond public health interventions, and there remains a desperate need for rapid development of antiviral therapeutics. A potential route to candidate antivirals is through repurposing of already approved drugs (for reviews; (2–4) and for examples;(5–8). We have previously screened a library of FDA approved drugs for antiviral activity against two other highly pathogenic human coronaviruses, SARS-CoV and Middle East respiratory syndrome coronavirus (MERS-CoV)(6). We found 27 drugs that inhibited replication of both of these coronaviruses, suggesting that they may have broad anti-coronaviral activity. One of the hits from this work was imatinib with which we subsequently determined the mechanism of action by demonstrating this drug inhibits fusion of coronaviruses with cellular membranes, thus blocking entry (9, 10).

Here, we present our investigation of 20 priority compounds from our previous screening to test if they can also inhibit SARS-CoV-2. Since these compounds are already approved for use in humans, they make ideal candidates for drug repurposing and rapid development as antiviral therapeutics. Our work found that 17 of the 20 of the drugs that inhibited SARS-CoV and MERS-CoV could also inhibit SARS-CoV-2, with similar IC50 values. We further assessed a subset of these drugs for their effects on SARS-CoV-2 RNA and infectious virus production and found all to have inhibitory activity. Our screening based on cytopathic effect therefore appears a favorable approach to find drugs capable of inhibiting production of infectious virus. Currently there are no established small animal model systems for SARS-CoV-2. However, there is a well-established mouse-adapted system for SARS-CoV (MA15 strain(11)) and we present data here assessing the *in vivo* efficacy of chloroquine (CQ) and chlorpromazine (CPZ) against SARS-CoV. We found that drug treatment does not inhibit virus replication in mouse lungs, but significantly improves clinical outcome. Based on both of these drugs inhibiting SARS-CoV-2 infection *in vitro* and providing protection *in vivo* against SARS-CoV clinical disease we believe they may be beneficial for SARS-CoV-2 therapy, but require further study in clinical contexts.

## Materials and Methods

### Cell lines and virus

Vero E6 cells (ATCC# CRL 1586) were cultured in DMEM (Quality Biological), supplemented with 10% (v/v) fetal bovine serum (Sigma), 1% (v/v) penicillin/streptomycin (Gemini Bio-products) and 1% (v/v) L-glutamine (2 mM final concentration, Gibco). Cells were maintained at 37°C and 5% CO_2_. Samples of SARS-CoV-2 were obtained from the CDC following isolation from a patient in Washington State (WA-1 strain - BEI #NR-52281). Stocks were prepared by infection of Vero E6 cells for two days when CPE was starting to be visible. Media were collected and clarified by centrifugation prior to being aliquoted for storage at −80°C. Titer of stock was determined by plaque assay using Vero E6 cells as described previously (12). All work with infectious virus was performed in a Biosafety Level 3 laboratory and approved by our Institutional Biosafety Committee. SARS-CoV stock was prepared as previously described (13). SARS-CoV spike (S) pseudotype viruses were produced as previously described (9).

### Drug testing

All drug screens were performed with Vero E6 cells. Cells were plated in opaque 96 well plates one day prior to infection. Drug stocks were made in either DMSO, water or methanol. Drugs were diluted from stock to 50 μM and an 8-point 1:2 dilution series made. Cells were pre-treated with drug for 2 hour (h) at 37°C/5% CO_2_ prior to infection at MOI 0.01 or 0.004. Vehicle controls were used on every plate, and all treatments were performed in triplicate for each screen. In addition to plates that were infected, parallel plates were left uninfected to monitor cytotoxicity of drug alone. Three independent screens with this set-up were performed. Cells were incubated at 37°C/5% CO_2_ for 3 days before performing CellTiter-Glo (CTG) assays as per the manufacturer’s instruction (Promega). Luminescence was read using a Molecular Devices Spectramax L plate reader. Fluphenazine dihydrochloride, benztropine mesylate, amodiaquine hydrochloride, amodiaquine dihydrochloride dihydrate, thiethylperazine maleate, mefloquine hydrochloride, triparanol, terconazole vetranal, anisomycin, fluspirilene, clomipramine hydrochloride, hydroxychloroquine sulfate, promethazine hydrochloride, emetine dihydrochloride hydrate and chloroquine phosphate were all purchased from Sigma. Chlorpromazine hydrochloride, toremifene citrate, tamoxifen citrate, gemcitabine hydrochloride and imatinib mesylate were all purchased from Fisher Scientific.

### Data analysis

Cytotoxicity (%TOX) data was normalized according to cell-only uninfected (cell only) controls and CTG-media-only (blank) controls:

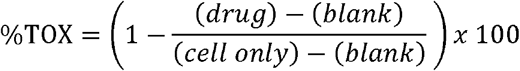

Inhibition (%Inhibit) data was normalized according to cell only and the activity of the vehicle controls:

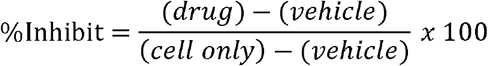

Nonlinear regression analysis was performed on the normalized %inhibit and %TOX data and IC50s and CC50s were calculated from fitted curves (log [agonist] versus response - variable slope [four parameters]) (GraphPad Software, LaJolla, CA), as described previously (14). Drug dilution points in a given run were excluded from IC50 analysis if the average cytotoxicity was greater than 30% (arbitrary cutoff) across the 3 cytotoxicity replicates for that screen. IC50 or CC50 values extrapolated outside the drug dilution range tested were reported as greater than 50μM or less than 0.39μM. Selectivity indexes (SI) were also calculated by dividing the CC50 by the IC50.

### Viral infection

To further analyse candidate drugs, Vero E6 cells were grown in 24 well plate format for 24 h prior to infection. As with the drug screens, cells were pre-treated with drug at a range of concentrations, or vehicle control for 2 h. Cells were then infected with SARS-CoV-2 at MOI 0.1 for 24 hour (h). Supernatant was collected, centrifuged in a table-top centrifuge for 3 minutes (min) at max speed and stored at −80°C. After a wash in PBS, infected cells were collected in TRIzol (Ambion) for RNA analysis (described below). Supernatant was used to titer viral production by TCID_50_ assay (12).

### RNA extraction and qRT-PCR

RNA was extracted from TRIzol samples using Direct-zol RNA miniprep kit (Zymo Research) as per the manufacturer’s instructions. RNA was converted to cDNA using RevertAid RT Kit (Thermo Scientific), with 12 μl of extracted RNA per reaction. For qRT-PCR, 2 μl of cDNA reaction product was mixed with PowerUp SYBR Green Master Mix (Applied Biosystems) and WHO/Corman primers targeting N and RdRp: N FWD 5’-CACATTGGCACCCGCAATC-3’, N REV 5’-GAGGAACGAGAAGAGGCTTG-3’, RdRp FWD 5’GTGARATGGTCATGTGTGGCGG-3’, RdRp REV 5’-CARATGTTAAASACACTATTAGCATA-3’. The qRT-PCR reactions were performed with a QuantStudio 5 (Applied Biosystems). To normalize loading, 18S RNA was used as a control, assessed with TaqMan Gene Expression Assays (Applied Biosystems) and TaqMan Fast Advanced Master Mix. Fold change between drug treated and vehicle control was determined by calculating ΔΔCT after normalization to the endogenous control of 18S.

### Pseudovirus fusion/entry assay

The pseudovirion (PV) entry assay was performed as described (9, 15). Briefly, 2 x 10^4^ BSC1 cells per well were in 96-well plates for 24 h, after which time cells were pre-treated with drug (1 h) and infected with PV (3 h). Media was removed and cells were washed with loading buffer (47 ml clear DMEM, 5 mM Probenecid, 2 mM L-glutamine, 25 mM HEPES, 200 nM bafilomycin, 5 μM E64D) and incubated for 1 h in CCF2 solution (LB, CCF2-AM, Solution B [CCF2-AM kit K1032] Thermo Fisher) in the dark. Cells were washed once with loading buffer and incubated from 6 h to overnight with 10% FBS in loading buffer. Percentage CCF2 cleavage was assessed by flow cytometry on the LSRII (Beckton Dickinson) in the flow cytometry core facility at the University of Maryland, Baltimore. Data were analyzed using FlowJo.

### Mouse infections

All infections were performed in an animal biosafety level 3 facility at the University of Maryland, Baltimore, using appropriate practices, including a HEPA-filtered bCON caging system, HEPA-filtered powered air-purifying respirators (PAPRs), and Tyvek suiting. All animals were grown to 10 weeks of age prior to use in experiments. The animals were anesthetized using a mixture of xylazine (0.38 mg/mouse) and ketamine (1.3 mg/mouse) in a 50 μl total volume by intraperitoneal injection. The mice were inoculated intranasally with 50 μl of either PBS or 2.5 x 10^3^ PFU of rMA15 SARS-CoV (11) after which all animals were monitored daily for weight loss. Mice were euthanized at day 4 post-infection, and lung tissue was harvested for further analysis. All animals were housed and used in accordance with the University of Maryland, Baltimore, Institutional Animal Care and Use Committee guidelines.

### Plaque assay

Vero cells were seeded in 35 mm dishes with 5 x 10^5^ cells per dish 24 h prior to infection. Supernatants from homogenized were serially diluted 10^−1^ through 10^−6^ in serum-free (SF) media. Cells were washed with SF media, 200 μl of diluted virus was added to each well and adsorption was allowed to proceed for 1 h at 37°C with gentle rocking every 10 min. 2X DMEM and 1.6% agarose were mixed 1:1. Cells were washed with SF media, 2 ml DMEM-agarose was added to each well, and cells were incubated for 72 h at 37°C, after which time plaques were read.

## Results

### Screening FDA approved compounds for anti-SARS-CoV-2 activity

Previously, we performed a large-scale drug screen on 290 FDA approved compounds to investigate which may have antiviral activity against SARS-CoV and MERS-CoV (6). With the emergence of SARS-CoV-2, we prioritized testing 20 of the 27 hits that were determined to inhibit both of the previously tested coronaviruses for antiviral activity against the novel virus. The list of tested compounds is shown in Table 1. Our screening started at 50 μM and used an 8-point, 1:2 dilution series with infections being performed at either MOI 0.01 or 0.004. CellTiter-Glo (CTG) assays were performed 3 days post-infection to determine relative cell viability between drug and vehicle control treated cells. Uninfected samples were used to measure the cytotoxicity of drug alone. From the relative luminescence data of the CTG assay, percent inhibition (of cell death caused by viral infection) could be measured and plotted along with the percent cytotoxicity of drug alone. Fig. 1 shows these plotted graphs from one representative of three independent screens at MOI 0.01. For those drugs demonstrating a cell toxicity rate lower than 30%, we were able to calculate IC50 values at both MOI from these graphs for 17 of the 20 drugs which is summarized in Table 1.

**Figure 1.**
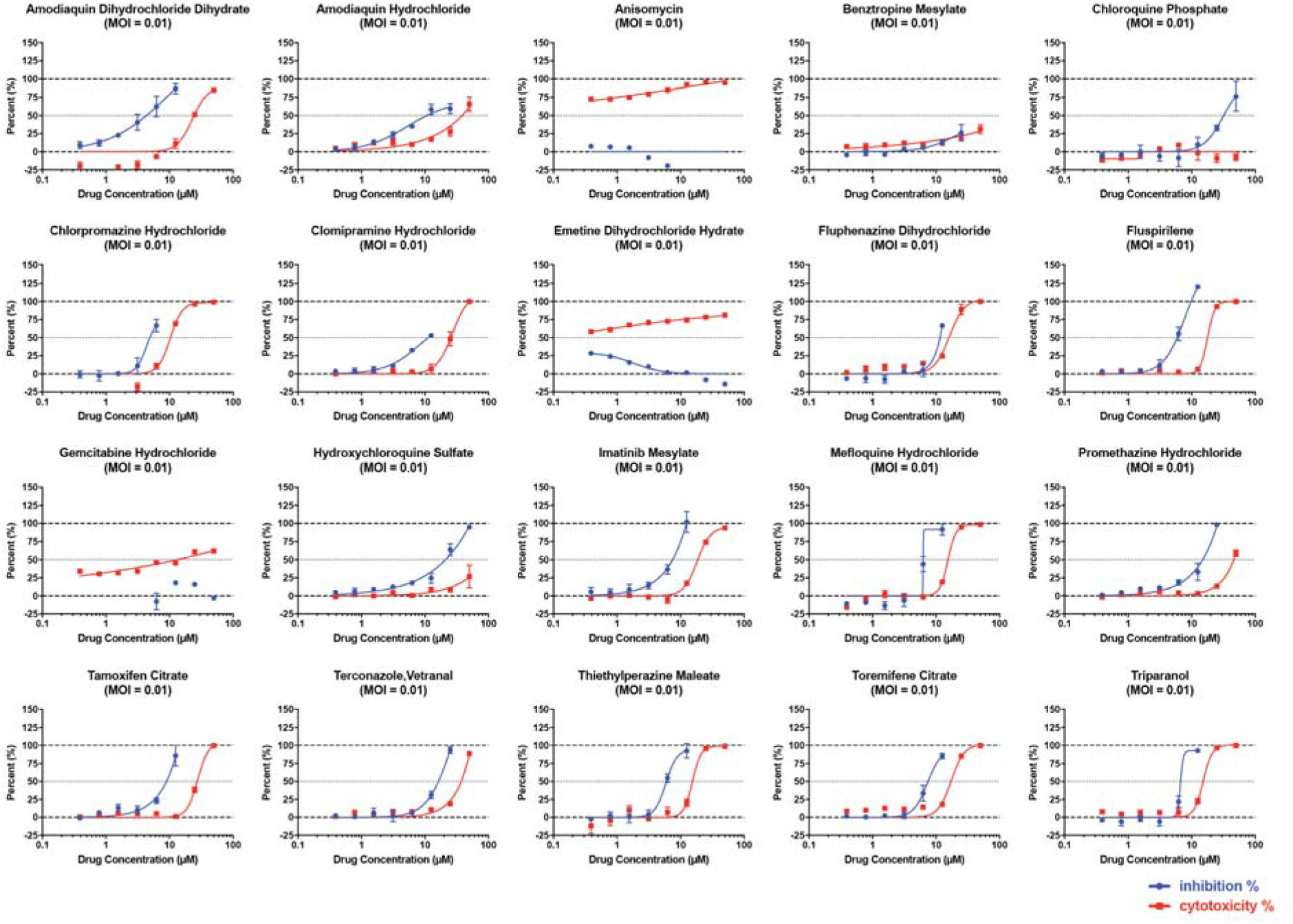
Percentage inhibition and percentage cytotoxicity graphs from drug screens starting at 50 μM using an 8-point, 1:2 dilution series. Results from one representative drug screen of three showing percentage inhibition and cytotoxicity for each of the tested drugs. Triplicate wells of cells were pre-treated with the indicated drug for 2 hours prior to infection with SARS-CoV-2 at MOI 0.01. Cells were incubated for 72 hours prior to performing CellTiter-Glo assays to assess cytopathic effect. Data are scored as percentage inhibition of relative cell viability for drug treated versus vehicle control. Data are the mean percentages with error bars displaying standard deviation between the triplicate wells.

**Table 1.**
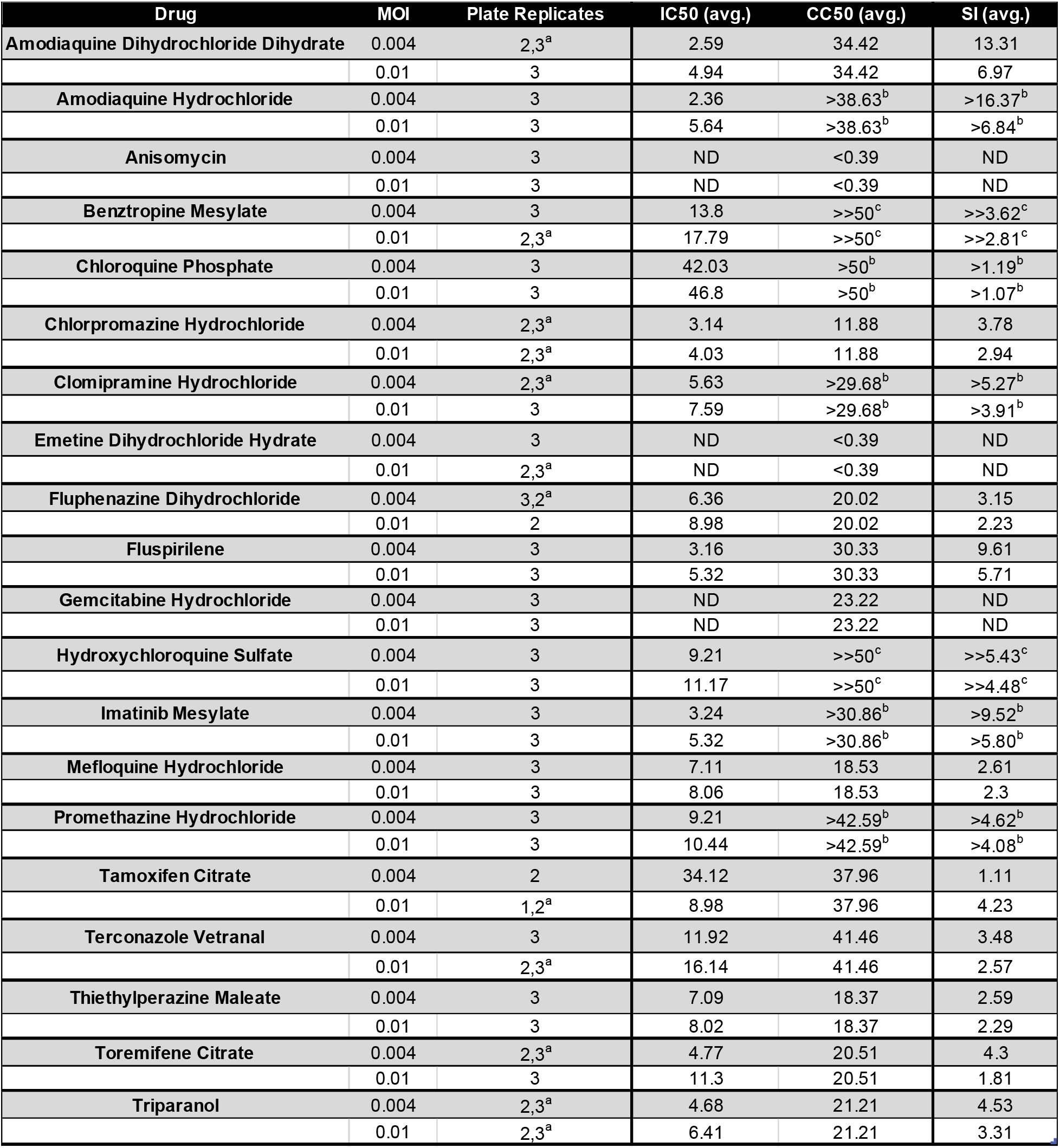
IC50 and CC50 values for 20 FDA approved drugs against SARS-CoV-2. Abbreviations: MOI (multiplicity of infection), IC50 (half maximal inhibitory concentration), CC50 (half maximal cytotoxic concentration), avg. (average), ND (not determined). A – Run totals listed as IC50, CC50 B – At least one CC50 could be extrapolated from the curve fit suggesting toxicity and SI are slightly higher than listed C – No CC50 could be extrapolated from the curve fit suggesting toxicity and SI are much higher than listed.

### Drug screen validation

In order to validate our screening process as a means to identify compounds with antiviral effect we decided to follow up with a subset of drugs. Chloroquine (CQ) has become the source of much interest as a potential treatment for COVID19 (16), as such, we further investigated hydroxychloroquine (HCQ) and CQ as both were present in our screen (Table 1). Vero E6 cells were plated and pre-treated with drug for 2 h prior to infection with SARS-CoV-2 at MOI 0.1. Supernatant was collected 24 h post-infection to determine titer of virus by TCID_50_ assay and cells were collected in TRIzol to assess production of viral mRNA. Treatment with both drugs caused a significant reduction in viral mRNA levels, especially at higher concentrations, without drug induced cytotoxicity (Fig. 1 and Table 1). There was a significant decrease in relative expression levels of both RdRp and N mRNA across the range of concentrations used (Fig. 2A-D). Along with causing a reduction in viral mRNA, treatment with both drugs caused a significant reduction in viral replication (Fig. 2E and 2F). SARS-CoV-2 production was more sensitive to HCQ than CQ with larger inhibition seen at the same concentration of treatment, which is in agreement with HCQ having a lower IC50 in our cell viability assay (Table 1). We also performed a time of addition assay with the highest concentration of HCQ to investigate whether SARS-CoV-2 entry was the point of inhibition of this compound (Fig. 2G). Interestingly, while the addition of HCQ at 2h post-infection did have some reduction in inhibitory activity there was not a complete loss, suggesting that HCQ treatment may impact other stages of the viral life cycle than just entry.

**Figure 2.**
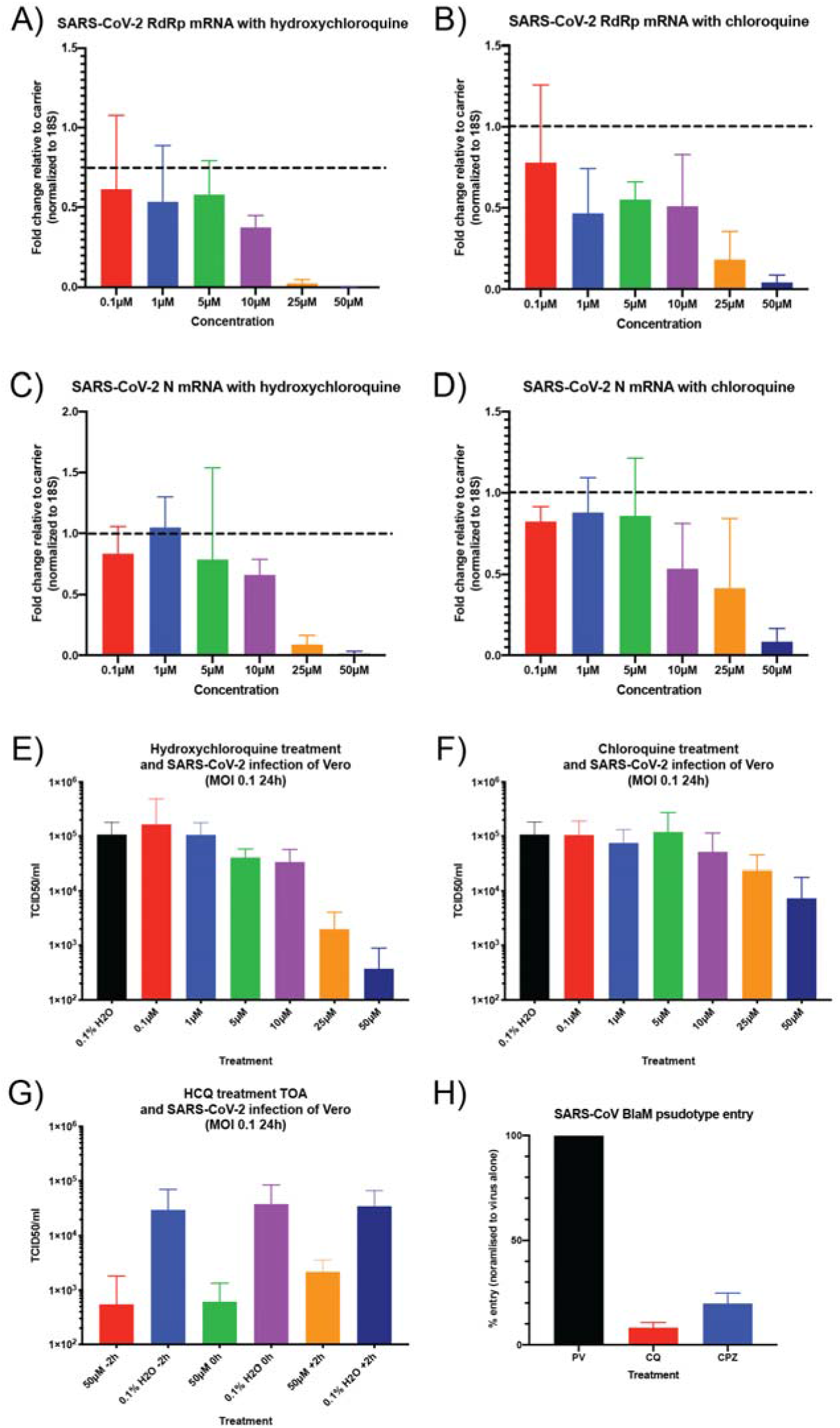
Hydroxychloroquine and chloroquine inhibit production of SARS-CoV-2 N and RdRp mRNA. Vero cells were pre-treated with hydroxychloroquine sulfate (A, C and E) or chloroquine phosphate (B, D and F) at the indicated concentration (or 0.1% water as vehicle control) for 2 h prior to infection with SARS-CoV-2 (WA-1 strain) at MOI 0.1. 24 h post-infection cells were collected in TRIzol. RNA was extracted from TRIzol sample and qRT-PCR was performed for viral RdRp (A and B) or N (C and D) mRNA using WHO primers. RNA levels were normalized with 18S RNA and fold change for drug treated to vehicle control was calculated (dotted line to denote a fold change of 1 which is no change over control). Data are from 3 independent infections performed on triplicate wells, the fold change was calculated in each independent experiment and the mean fold change is plotted with error bars displaying standard deviation. Along with TRIzol samples for RNA supernatant was collected from cells and used for TCID_50_ assays to determine infectious virus production following treatment with HCQ (E) or CQ (F) Data are from 3 independent infections performed on triplicate wells with the TCID_50_/ml being averaged across all wells. Error bars are the standard deviation. G) Cells were treated with 50 μM HCQ or 0.1% water as control. Drug was either added 2 h prior to infection, at the time of infection or 2 h after infection with MOI 0.1 SARS-CoV-2. After 24 h infection, supernatant was collected and used for TCID_50_ assays to determine infectious virus production. Data are from 3 independent infections performed on triplicate wells with the TCID_50_/ml being averaged across all wells. Error bars are the standard deviation. H) SARS-CoV spike psuedoviruses (PV) were used for infection of BSC1 cells. The cells were treated with 10 μM of CQ or CPZ for 1 h prior to infection with PV for 3 h. The PV carry BlaM and cells were loaded with CCF2 to monitor cleavage and shift in fluorescence output for evidence of S-mediated entry into cells. Data are normalised to PV alone and are from 3 independent experiments with error bars representing standard deviation.

We have previously used a β-lactamase-Vpr chimeric protein (Vpr-BlaM) pseudotype system to demonstrate that imatinib (a drug also seen to inhibit SARS-CoV-2 [Table 1 and Fig. 1]) inhibits SARS-CoV and MERS-CoV spike-mediated entry (9). We used this system to more directly investigate whether CQ could inhibit viral entry mediated by coronavirus spike, and additionally included chlorpromazine (CPZ) as this is known to inhibit clathrin-mediated endocytosis (17) and was also part of our drug screening (Table 1 and Fig. 2). In this assay, when the pseudovirus fuses with a cellular membrane, BlaM is released into the cytoplasm of the infected cell. BlaM cleaves cytoplasmic loaded CCF2 to change its emission spectrum from 520 nm (green) to ~450 nm (blue), which can be quantified using flow cytometry.

Cells were treated with CQ or CPZ for 1 h before infection with BlaM-containing SARS-S pseudovirions (PV). Cells were then analyzed by flow cytometry to quantify the cleavage of CCF2. In mock treated cells infected with SARS-S PV there was a shift in the CCF2 emission spectrum indicating release of BlaM to the cytosol, and that spike-mediated fusion with cellular had occurred. Upon treatment with CQ or CPZ there was a greater than 90% reduction in CCF2 cleavage caused by SARS-S PV (Fig. 2H). These data demonstrate that both drugs inhibit SARS-CoV spike-mediated fusion with cellular membranes. These pseudotype assays suggest that the inhibition of coronavirus replication caused by CQ and CPZ is at the stage of entry to cells but combined with the time of addition assays (Fig. 2G), there is a suggestion that later stages may also be impacted.

HCQ and CQ are used as anti-malarial drugs and are in the class of aminoquinolines which are hemozoin inhibitors, similarly to 4-methanolquinolines. Interestingly, from our drug screening, three other hemozoin inhibitors were identified: amodiaquine dihydrochloride dihydrate, amodiaquine hydrochloride and mefloquine. We therefore decided to directly test these drugs for antiviral activity against SARS-CoV-2. We directly tested CPZ against SARS-CoV-2 having seen that it could inhibit SARS-CoV S-mediated entry to cells (Fig. 2H). We also included imatinib since we have previously shown that this can inhibit entry of both SARS-CoV and MERS-CoV (9) and was a hit against SARS-CoV-2 (Fig. 1 and Table 1). Again, cells were pre-treated with drugs at the indicated concentrations and infected with SARS-CoV-2 at MOI 0.1 for 24h, after which supernatant samples were collected. As can be seen in Fig. 3, at the highest concentrations of all drugs there is significant inhibition of SARS-CoV-2 infection. All five drugs showed very strong inhibition at 20 μM.

**Figure 3.**
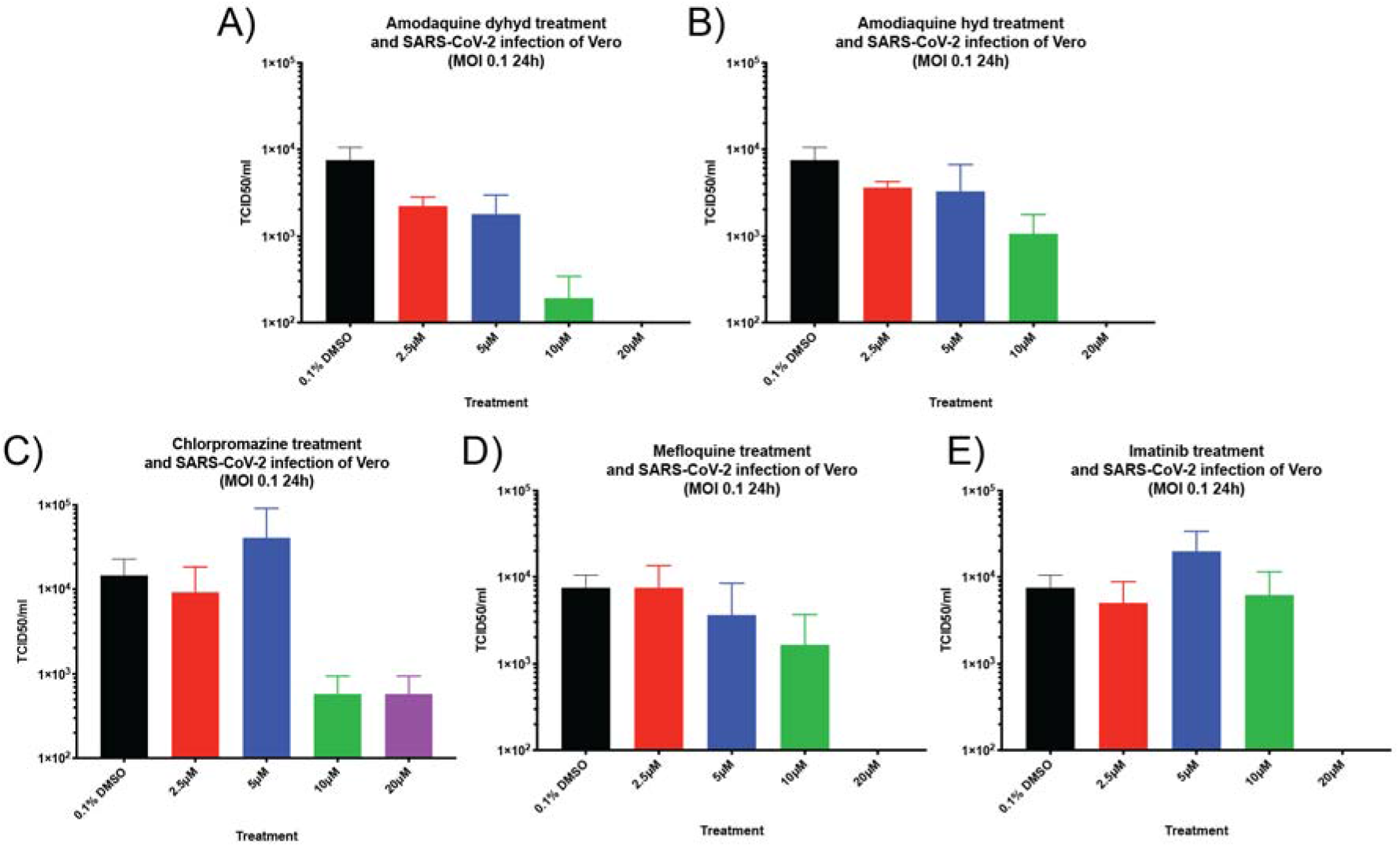
Antiviral activity of additional FDA approved compounds against SARS-CoV-2. Other drugs that showed antiviral activity in our initial CellTiter-Glo screening were tested for inhibition of productive virus infection. Cells were treated with the indicated concentrations of A) amodiaquine dihydrochloride dihydrate, B) amodiaquine hydrochloride, C) chlorpromazine, D) mefloquine and E) imatinib for 2 h prior to infection with SARS-CoV-2 at MOI 0.1 for 24 h. Supernatant was collected and used for TCID_50_ assay to quantify infectious virus production. Data are from a representative experiment of four performed on triplicate wells. Data are the mean TCID_50_/ml with error bars being standard deviation.

Overall, the data from Fig. 2 and Fig. 3 indicate that there are various FDA approved drugs that have broad-spectrum anti-coronavirus activity *in vitro* and that our initial screening based on cytopathic effect is a good method to identify compounds with antiviral activity.

### Chloroquine and chlorpromazine do not inhibit SARS-CoV (MA15) replication in mouse lungs, but significantly reduces weight loss and clinical signs

CQ and CPZ treatment displayed significant inhibition of coronavirus replication *in vitro*, with our data suggesting entry is inhibited. We therefore decided to investigate whether these drugs were efficacious *in vivo* using SARS-CoV strain, MA15 in BALB/c mice. This model displays ~15-20% weight loss by 4 days post infection (dpi), occasionally resulting in death. We tested whether prophylactically administered CQ or CPZ could protect mice from severe MA15 infection. Mice were injected intraperitoneally with either water, 0.8 mg CQ, 1.6 mg CQ, 20 μg CPZ, 100 μg CPZ or 200 μg CPZ at day −1 of infection and then were dosed every day through the 4 days of infection. On day 0, mice were intranasally infected with 2.5 x 10^3^ pfu of SARS-CoV (MA15) or PBS as control. Weight loss was measured as a correlate of disease and mice were euthanized at 4 dpi for analysis.

PBS inoculated mice showed no weight loss or clinical signs of disease when treated with either water, CQ or CPZ over the experiment time course indicating drug treatment did not adversely affect morbidity (Fig. 4A and 4D). Mice that were infected with MA15 and treated with water lost ~15% of their starting body weight over 4 days and had significant clinical signs of disease including ruffled fur, labored breathing and lethargy (Fig. 4A and 4D). Mice that were treated with 0.8 mg of CQ each day, displayed similar weight loss as the water control through the first 3 days of infection, however by 4 dpi the weight loss was halted in the drug treated mice (Fig. 4A). Mice that were treated with 1.6 mg CQ per day showed markedly reduced weight loss compared to the water control (Fig. 4A). Pathological analysis was also performed on H&E stained sections. Mice infected with MA15 and treated with water displayed significant inflammation and denuding bronchiolitis suggesting severe disease (Fig. 4B). By contrast, 0.8 mg CQ dose group had moderate inflammation that was reduced compared to control and the 1.6 mg dose group had minimal lung pathology (Fig. 4B). Interestingly, even though CQ treatment appeared to protect against weight loss and inflammation in the lungs, the viral titer was equivalent between drug treated and vehicle control mice (Fig. 4C).

**Figure 4.**
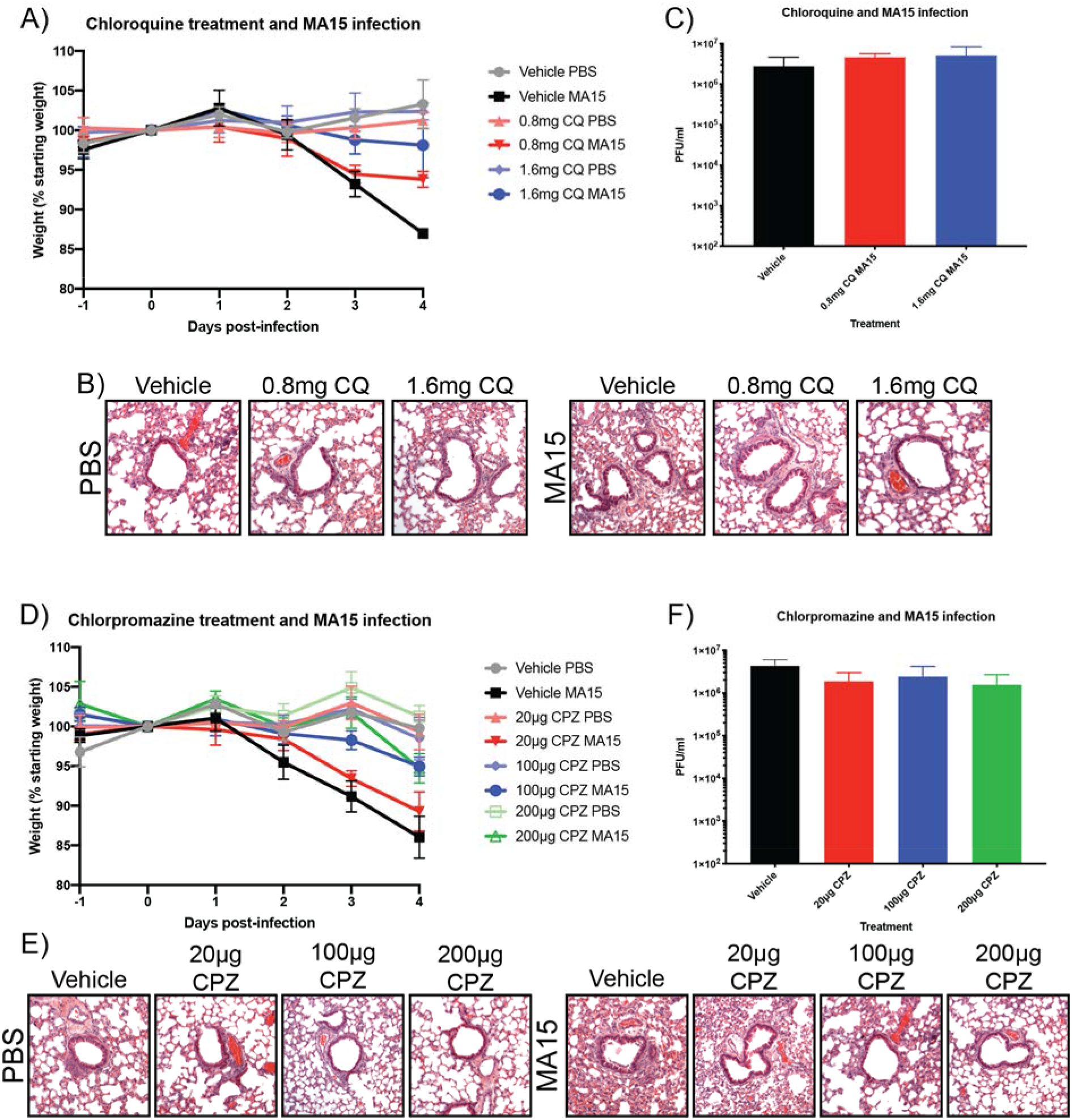
CQ and CPZ are protective against SARS-CoV (MA15) infection *in vivo*. Mice were treated with CQ or CPZ 1 day prior to infection with SARS-CoV (MA15) and dosed with each drug across the 4 day infection time course. Water was used as the vehicle control for both drugs and PBS was used as a control for uninfected mice. A) Weight loss of mice treated with CQ at two different dose levels (0.8 mg and 1.6 mg) over the 4 day infection. Data are presented as relative weight loss compared to the mouse weight on day 0. In each treatment group there were 5 mice and the data are mean average and standard deviation. B) At day 4, mice were euthanized and lung sections were used for H&E staining. C) In addition to collecting lungs for section staining, there was also collection to determine titer of virus by plaque assay. D) Weight loss of mice treated with CPZ at three different doses (20 μg, 100 μg, and 200 μg) with the same experimental set up as in A. E and F) As B and C but for CPZ treated mice.

Similar to the CQ results, CPZ treatment reduced weight loss in mice infected with MA15 at 100 μg and 200 μg, but the 20 μg treatment group were equivalent to vehicle control (Fig. 4D) and the H&E sections showed protection against inflammation and denuding bronchiolitis at the higher doses (Fig. 4E). Again, as with CQ treatment, even though there were reduced signs of infection with CPZ treatment, there was no difference in MA15 titer in the mouse lungs (Fig. 4F). Overall these data indicate that even though CQ and CPZ treatment do not inhibit viral replication in the lungs, both can protect mice from signs of disease following SARS-CoV (MA15) infection.

## Discussion

The SARS-CoV-2 pandemic has demonstrated the desperate need for antiviral drugs. Since the emergence of SARS-CoV in 2002, research has uncovered many details of coronavirus biology and pathogenesis, however there are currently no approved therapeutics against this emerging virus family. Whether being used for treating SARS-CoV-2 in this current pandemic or the next unknown viral pathogen in the future, we must attempt to develop and validated antiviral drugs that are ready to be used at the first signs of an outbreak. Many FDA approved drugs have been found to have antiviral activity in addition to their approved use (e.g; (5–8)), and since these are extensively used in humans for other conditions, they could be streamlined for rapid approval and repositioning as antivirals. In our previous work, 290 FDA approved drugs were screened for antiviral activity and 27 were found to inhibit both SARS-CoV and MERS-CoV (6). We prioritized testing these for antiviral activity against SARS-CoV-2 since they displayed broad-spectrum antiviral activity. From multiple independent screens performed with two MOI, we found that 17 of our 20 tested priority compounds display significant antiviral activity at non-cytotoxic concentrations. Many of the compounds have IC50 values under 10 μM and these will be the source of follow up testing on additional cell lines and in mouse models of SARS-CoV-2. We further investigated seven of the hits to directly test if they inhibited SARS-CoV-2 replication. We performed follow-up experiments with HCQ, CQ, amodiaquine and mefloquine because chloroquine has garnered much interest as a potential treatment for COVID19 (16) and the others are similarly used as anti-malarial compounds (18). In addition, we have previously demonstrated that imatinib is an inhibitor of SARS-CoV, MERS-CoV and infectious bronchitis virus entry to cells (9, 10) so included that here as the mechanism of coronavirus inhibition is understood. Finally, CPZ inhibits clathrin function in cells (17) so can disrupt infection by many viruses that require clathrin-mediated endocytosis and was therefore also chosen for further analysis. Treatment of cells with all these drugs showed inhibition of infectious viral particle production (measured by TCID_50_ assay) at non-cytotoxic levels.

Having demonstrated that HCQ, CQ and CPZ can inhibit cytopathic effect, mRNA synthesis and infectious viral particle production of SARS-CoV-2, we used a previously published system of SARS-CoV pseudotype viruses carrying Vpr-BlaM to investigate whether CQ and CPZ inhibit coronavirus spike-mediated entry to better define mechanism of action. We have previously used this system to define imatinib as an entry inhibitor of these viruses (9) and found similar results for CQ and CPZ, thus better defining their mechanism of antiviral activity.

Finally, we investigated the efficacy of CQ and CPZ with an *in vivo* model using SARS-CoV MA15. There is currently a lack of an established mouse model for SARS-CoV-2 so we used the mouse adapted SARS-CoV (MA15) strain as a surrogate to assess the *in vivo* efficacy of these drugs against a closely related coronavirus. We are of the opinion that this is a good model since both viruses use ACE2 as a receptor (19–22) and therefore have a similar cellular tropism which is important since both of these compounds appear to inhibit viral entry. Prophylactic dosing in MA15 infection experiments demonstrated that, in contrast to the *in vitro* antiviral activity, CQ and CPZ did not inhibit viral replication in mouse lungs based on viral titer recovered from lungs at 4 dpi. However, both drugs resulted in reduced weight loss and improved clinical outcome, with the higher dose giving greater protection. Along with being an anti-malarial, CQ is used in humans for the treatment of systemic lupus erythematosus and rheumatoid arthritis because of anti-inflammatory properties and has effects on antigen presentation (23–25). We speculate that these properties may have a role in the protection we observe *in vivo* since much of the pathology from SARS-CoV is a consequence of immunopathology during infection (in mice; (26), in non-human primates (27) and for a detail review (28)). These results suggest that CQ alone may not be a viable therapeutic but may be beneficial for treatment of SARS-CoV-2 in combination with more directly acting antivirals such as remdesivir (29–31).

The development of antiviral drugs for emerging coronaviruses is a global priority. In the middle of the COVID19 pandemic, we must identify rapidly accessible therapeutics that are validated in both *in vitro* and *in vivo* models. FDA approved drugs being assessed for repurposing and other experimental drugs in development must be properly validated in animal studies to best assess their potential utility in people. We have presented here a list of FDA approved drugs that are effective *in vitro* against SARS-CoV-2 as well as being effective against SARS-CoV and MERS-CoV (6). Moreover, we have demonstrated that two of these, CQ and CPZ, can protect mice from severe clinical disease from SARS-CoV. Future research will be aimed at testing these compounds in SARS-CoV-2 animal models to further assess their potential utility for human treatment.

## Acknowledgments

We kindly thank Emergent BioSolutions for financial support to perform these experiments. We also kindly thank Julie Dyall for helpful discussions regarding data analysis.

## Notes

### Competing Interest Statement

The authors have declared no competing interest.

### Summary of Updates

The previous manuscript had the data that is in Fig 1 and had 2 independent repeats of the RNA and TCID50 experiments in Fig. 2. Those experiments in Fig. 2 now have three independent repeats. All other data in Fig. 2, 3 and 4 is new data added to the updated manuscript. As part of the data that has been added, two additional authors have been added in CMC and JMS.

## References

1. Zhu N, Zhang D, Wang W, Li X, Yang B, Song J, Zhao X, Huang B, Shi W, Lu R, Niu P, Zhan F, Ma X, Wang D, Xu W, Wu G, Gao GF, Tan W. 2020. A novel coronavirus from patients with pneumonia in China, 2019. N Engl J Med 382:727–733.

2. Sisk JM, Frieman MB. 2016. Screening of FDA-Approved Drugs for Treatment of Emerging Pathogens. ACS Infect Dis. American Chemical Society.

3. Pushpakom S, Iorio F, Eyers PA, Escott KJ, Hopper S, Wells A, Doig A, Guilliams T, Latimer J, McNamee C, Norris A, Sanseau P, Cavalla D, Pirmohamed M. 2018. Drug repurposing: Progress, challenges and recommendations. Nat Rev Drug Discov.

4. Mercorelli B, Palù G, Loregian A. 2018. Drug Repurposing for Viral Infectious Diseases: How Far Are We? Trends Microbiol 26:865–876.

5. Madrid PB, Chopra S, Manger ID, Gilfillan L, Keepers TR, Shurtleff AC, Green CE, Iyer L V., Dilks HH, Davey RA, Kolokoltsov AA, Carrion R, Patterson JL, Bavari S, Panchal RG, Warren TK, Wells JB, Moos WH, Burke RLL, Tanga MJ. 2013. A Systematic Screen of FDA-Approved Drugs for Inhibitors of Biological Threat Agents. PLoS One 8.

6. Dyall J, Coleman CM, Hart BJ, Venkataraman T, Holbrook MR, Kindrachuk J, Johnson RF, Olinger GG, Jahrling PB, Laidlaw M, Johansen LM, Lear-rooney CM, Glass PJ, Hensley LE, Frieman B. 2014. Repurposing of Clinically Developed Drugs for Treatment of Middle East Respiratory Syndrome Coronavirus Infection. Antimicrob Agents Chemother 58:4885–4893.

7. Madrid PB, Panchal RG, Warren TK, Shurtleff AC, Endsley AN, Green CE, Kolokoltsov A, Davey R, Manger ID, Gilfillan L, Bavari S, Tanga MJ. 2016. Evaluation of Ebola Virus Inhibitors for Drug Repurposing. ACS Infect Dis 1:317–326.

8. Xu M, Lee EM, Wen Z, Cheng Y, Huang WK, Qian X, Tcw J, Kouznetsova J, Ogden SC, Hammack C, Jacob F, Nguyen HN, Itkin M, Hanna C, Shinn P, Allen C, Michael SG, Simeonov A, Huang W, Christian KM, Goate A, Brennand KJ, Huang R, Xia M, Ming GL, Zheng W, Song H, Tang H. 2016. Identification of small-molecule inhibitors of Zika virus infection and induced neural cell death via a drug repurposing screen. Nat Med 22:1101–1107.

9. Coleman CM, Sisk JM, Mingo RM, Nelson EA, White JM, Frieman MB. 2016. Abl Kinase Inhibitors Are Potent Inhibitors of SARS-CoV and MERS-CoV Fusion. J Virol 90:8924–8933.

10. Sisk JM, Frieman MB, Machamer CE. 2018. Coronavirus S protein-induced fusion is blocked prior to hemifusion by Abl kinase inhibitors. J Gen Virol 1–12.

11. Roberts A, Deming D, Paddock CD, Cheng A, Yount B, Vogel L, Herman BD, Sheahan T, Heise M, Genrich GL, Zaki SR, Baric R, Subbarao K. 2007. A mouse-adapted SARS-coronavirus causes disease and mortality in BALB/c mice. PLoS Pathog 3:0023–0037.

12. Coleman CM, Frieman MB. 2015. Growth and Quantification of MERS-CoV Infection. Curr Protoc Microbiol 37:15E.2.1–15E.2.9.

13. Frieman M, Yount B, Agnihothram S, Page C, Donaldson E, Roberts A, Vogel L, Woodruff B, Scorpio D, Subbarao K, Baric RS. 2012. Molecular Determinants of Severe Acute Respiratory Syndrome Coronavirus Pathogenesis and Virulence in Young and Aged Mouse Models of Human Disease. J Virol 86:884–897.

14. Dyall J, Johnson JC, Hart BJ, Postnikova E, Cong Y, Zhou H, Gerhardt DM, Michelotti J, Honko AN, Kern S, DeWald LE, O’Loughlin KG, Green CE, Mirsalis JC, Bennett RS, Olinger GG, Jahrling PB, Hensley LE. 2018. In Vitro and In Vivo Activity of Amiodarone Against Ebola Virus. J Infect Dis 218:S592–S596.

15. Mingo RM, Simmons JA, Shoemaker CJ, Nelson EA, Schornberg KL, D’Souza RS, Casanova JE, White JM. 2015. Ebola Virus and Severe Acute Respiratory Syndrome Coronavirus Display Late Cell Entry Kinetics: Evidence that Transport to NPC1 + Endolysosomes Is a Rate-Defining Step. J Virol 89:2931–2943.

16. Pastick KA, Okafor EC, Wang F, Lofgren SM, Skipper CP, Nicol MR, Pullen MF, Rajasingham R, Mcdonald EG, Lee TC, Schwartz IS, Kelly LE, Lother SA, Mitjà O, Letang E, Abassi M, Boulware DR. 2020. Review: Hydroxychloroquine and Chloroquine for Treatment of SARS-CoV-2 (COVID-19). Open Forum Infect Dis.

17. Wang LH, Rothberg KG, Anderson RGW. 1993. Mis-assembly of clathrin lattices on endosomes reveals a regulatory switch for coated pit formation. J Cell Biol 123:1107–1117.

18. Baird JK. 2005. Effectiveness of antimalarial drugs. N Engl J Med 352.

19. Li W, Moore MJ, Vasllieva N, Sui J, Wong SK, Berne MA, Somasundaran M, Sullivan JL, Luzuriaga K, Greeneugh TC, Choe H, Farzan M. 2003. Angiotensin-converting enzyme 2 is a functional receptor for the SARS coronavirus. Nature 426:450–454.

20. Zhou P, Yang X Lou, Wang XG, Hu B, Zhang L, Zhang W, Si HR, Zhu Y, Li B, Huang CL, Chen HD, Chen J, Luo Y, Guo H, Jiang R Di, Liu MQ, Chen Y, Shen XR, Wang X, Zheng XS, Zhao K, Chen QJ, Deng F, Liu LL, Yan B, Zhan FX, Wang YY, Xiao GF, Shi ZL. 2020. A pneumonia outbreak associated with a new coronavirus of probable bat origin. Nature 579:270–273.

21. Wan Y, Shang J, Graham R, Baric RS, Li F. 2020. Receptor Recognition by the Novel Coronavirus from Wuhan: an Analysis Based on Decade-Long Structural Studies of SARS Coronavirus. J Virol 94.

22. Hoffmann M, Kleine-Weber H, Schroeder S, Krüger N, Herrler T, Erichsen S, Schiergens TS, Herrler G, Wu NH, Nitsche A, Müller MA, Drosten C, Pöhlmann S. 2020. SARS-CoV-2 Cell Entry Depends on ACE2 and TMPRSS2 and Is Blocked by a Clinically Proven Protease Inhibitor. Cell 181:271–280.e8.

23. Ziegler HK, Unanue ER. 1982. Decrease in macrophage antigen catabolism caused by ammonia and chloroquine is associated with inhibition of antigen presentation to T cells. Proc Natl Acad Sci U S A 79:175–178.

24. Al-Bari MAA. 2015. Chloroquine analogues in drug discovery: new directions of uses, mechanisms of actions and toxic manifestations from malaria to multifarious diseases. J Antimicrob Chemother 70:1608–21.

25. Rainsford KD, Parke AL, Clifford-Rashotte M, Kean WF. 2015. Therapy and pharmacological properties of hydroxychloroquine and chloroquine in treatment of systemic lupus erythematosus, rheumatoid arthritis and related diseases. Inflammopharmacology 23:231–269.

26. Rockx B, Baas T, Zornetzer GA, Haagmans B, Sheahan T, Frieman M, Dyer MD, Teal TH, Proll S, van den Brand J, Baric R, Katze MG. 2009. Early Upregulation of Acute Respiratory Distress Syndrome-Associated Cytokines Promotes Lethal Disease in an Aged-Mouse Model of Severe Acute Respiratory Syndrome Coronavirus Infection. J Virol 83:7062–7074.

27. Smits SL, De Lang A, Van Den Brand JMA, Leijten LM, Van Ijcken WF, Eijkemans MJC, Van Amerongen G, Kuiken T, Andeweg AC, Osterhaus ADME, Haagmans BL. 2010. Exacerbated innate host response to SARS-CoV in aged non-human primates. PLoS Pathog 6.

28. Channappanavar R, Perlman S. 2017. Pathogenic human coronavirus infections: causes and consequences of cytokine storm and immunopathology. Semin Immunopathol 39:529–539.

29. Wang M, Cao R, Zhang L, Yang X, Liu J, Xu M, Shi Z, Hu Z, Zhong W, Xiao G. 2020. Remdesivir and chloroquine effectively inhibit the recently emerged novel coronavirus (2019-nCoV) in vitro. Cell Res 0.

30. Brown AJ, Won JJ, Graham RL, Dinnon KH, Sims AC, Feng JY, Cihlar T, Denison MR, Baric RS, Sheahan TP. 2019. Broad spectrum antiviral remdesivir inhibits human endemic and zoonotic deltacoronaviruses with a highly divergent RNA dependent RNA polymerase. Antiviral Res 169.

31. Sheahan TP, Sims AC, Graham RL, Menachery VD, Gralinski LE, Case JB, Leist SR, Pyrc K, Feng JY, Trantcheva I, Bannister R, Park Y, Babusis D, Clarke MO, MacKman RL, Spahn JE, Palmiotti CA, Siegel D, Ray AS, Cihlar T, Jordan R, Denison MR, Baric RS. 2017. Broad-spectrum antiviral GS-5734 inhibits both epidemic and zoonotic coronaviruses. Sci Transl Med 9.

